# Limited variation between SARS-CoV-2-infected individuals in domain specificity and relative potency of the antibody response against the spike glycoprotein

**DOI:** 10.1101/2021.08.04.455181

**Authors:** Hanora A. Van Ert, Dana W. Bohan, Kai Rogers, Mohammad Fili, Roberth A. Rojas Chávez, Enya Qing, Changze Han, Spencer Dempewolf, Guiping Hu, Nathan Schwery, Kristina Sevcik, Natalie Ruggio, Devlin Boyt, Michael A. Pentella, Tom Gallagher, J. Brooks Jackson, Anna E. Merrill, C. Michael Knudson, Grant D. Brown, Wendy Maury, Hillel Haim

## Abstract

The spike protein of SARS-CoV-2 is arranged as a trimer on the virus surface, composed of three S1 and three S2 subunits. Infected and vaccinated individuals generate antibodies against spike, which can neutralize the virus. Most antibodies target the receptor-binding domain (RBD) and N-terminal domain (NTD) of S1; however, antibodies against other regions of spike have also been isolated. The variation between infected individuals in domain specificity of the antibodies and in their relative neutralization efficacy is still poorly characterized. To this end, we tested serum and plasma samples from 85 COVID-19 convalescent subjects using 7 immunoassays that employ different domains, subunits and oligomeric forms of spike to capture the antibodies. Samples were also tested for their neutralization of pseudovirus containing SARS-CoV-2 spike and of replication-competent SARS-CoV-2. We observed strong correlations between the levels of NTD- and RBD-specific antibodies, with a fixed ratio of each type to all anti-spike antibodies. The relative potency of the response (defined as the measured neutralization efficacy relative to the total level of spike-targeting antibodies) also exhibited limited variation between subjects, and was not associated with the overall amount of anti-spike antibodies produced. Accordingly, the ability of immunoassays that use RBD, NTD and different forms of S1 or S1/S2 as capture antigens to estimate the neutralizing efficacy of convalescent samples was largely similar. These studies suggest that host-to-host variation in the polyclonal response elicited against SARS-CoV-2 spike is primarily limited to the quantity of antibodies generated rather than their domain specificity or relative neutralization potency.

**IMPORTANCE:** Infection by SARS-CoV-2 elicits antibodies against various domains of the spike protein, including the RBD, NTD and S2. Different infected individuals generate vastly different amounts of anti-spike antibodies. By contrast, as we show here, there is a remarkable similarity in the properties of the antibodies produced. Different individuals generate the same proportions of antibodies against each domain of the spike protein. Furthermore, the relationship between the amount of anti-spike antibodies produced and their neutralization efficacy of SARS-CoV-2 is highly conserved. Therefore, the observed variation in the neutralizing activity of the antibody response in COVID-19 convalescent subjects is caused by differences in the amounts of antibodies rather than their recognition properties or relative antiviral activity. These findings suggest that COVID-19 vaccine strategies that focus on enhancing the overall level of the antibodies will likely elicit a more uniformly efficacious protective response.

## INTRODUCTION

The spike protein on the surface of SARS-CoV-2 mediates fusion with target cells (1, 2). Spike is generated as a precursor that is cleaved by furin in the producer cells to generate S1 and S2 subunits (3). These subunits are non-covalently associated on the virus surface, where they form a trimer of heterodimers (4). Furin cleavage primes spike for further processing by the serine protease TMPRSS2 on the plasma membrane or the cysteine protease cathepsin L within the endosome (3, 5, 6). Spike is highly immunogenic in humans and, in infected and vaccinated individuals, readily elicits antibodies that play a critical role in protection (7, 8). Most neutralizing antibodies isolated to date target the receptor-binding domain (RBD) on the S1 subunit (9–15). In addition, multiple neutralizing antibodies that target the N-terminal domain (NTD) of S1 have been isolated (16–18). By contrast, neutralizing antibodies against the C-terminal domain (CTD) of S1 or against the S2 subunit are relatively rare (19, 20). The variation between individuals in the domain specificity of the anti-spike response and in the relative neutralization efficacy of the antibodies produced remains poorly explored.

To address this question, we quantified the binding specificity of anti-spike antibodies in 85 convalescent COVID-19 serum and plasma samples using capture antigens that represent different domains, subunits, and oligomeric forms of spike. A panel of 7 in-house and commercial immunoassays that quantify anti-spike antibodies was tested, as well as a nucleocapsid-based assay. Antibody content in the samples measured by these assays was compared with their neutralization efficacy of SARS-CoV-2. We observed that different subjects exhibit remarkably similar ratios of anti-RBD and anti-NTD antibodies relative to the total anti-spike antibodies. Interestingly, the relative potency of the convalescent samples (defined as the ratio between neutralization efficacy and the amount of anti-spike antibodies measured) was also similar in different individuals, and was not associated with the robustness of the response against spike. Our results demonstrate limited host-to-host variation in both spike domain specificity and in the relative potency of the antibody response elicited after SARS-CoV-2 infection. Variation between hosts in the polyclonal response generated is primarily limited to the quantity of the antibodies rather than the domains targeted or the efficacy of their neutralizing activity.

## RESULTS

### Strong correlations between results of immunoassays that apply different spike components as the capture antigen

To determine the target specificity of the SARS-CoV-2 antibody response, we analyzed serum and plasma samples collected from individuals who had recovered from COVID-19 (see all donor information in **Supp. Table S1**). Serum samples were obtained from individuals willing to donate convalescent plasma for the treatment of COVID-19 patients. In addition, we analyzed plasma samples from obstetric patients who had serologic evidence of COVID-19 infection; samples were collected during their hospitalization for delivery (21). All samples were collected between March 2020 and January 2021. None of the donors required hospitalization for COVID-19-related symptoms. For 68% of donors, the precise date of positivity for SARS-CoV-2 (by PCR analysis of a nasopharyngeal swab) was known; among these subjects, 82% of the serum or plasma samples were collected within 60 days of the positive PCR result (**Supp. Fig. S1A**). For each serum or plasma sample, we quantified the levels of SARS-CoV-2-specific antibodies using commercially-available and in-house-developed immunoassays that apply different domains, subunits or oligomeric forms of the spike protein as the capture antigen (**Table 1** and **Fig. 1A**). To mimic the native spike trimer on the virus surface, we used a cell-based enzyme linked immunosorbent assay (cbELISA) that measures antibodies against the full-length membrane-bound form of spike (22–24). For this purpose, we used human osteosarcoma (HOS) cells that express on their surface fusion-competent spike trimers by transfection with an expression plasmid that encodes the full-length protein. Samples were also tested by ELISAs, in which recombinant soluble dimeric forms of the RBD, NTD or the complete ectodomain of S1/S2 (designated Ecto) were used as the capture antigens. The Ecto protein was generated by abrogating the furin cleavage site at spike positions 682-685 (3). Binding of antibodies in serum or plasma to these antigens was measured using a secondary antibody specific for the human kappa light chain, which detects isotypes IgG, IgM and IgA. In addition, we tested the samples with commercial immunoassays that detect IgG against the S1 subunit (Ortho Vitros), S1/S2 subunits (DiaSorin Liaison IgG) and a trimeric soluble form of S1/S2 (DiaSorin TrimericS IgG). To quantify non-spike-targeting antibodies elicited against SARS-CoV-2, we used the Roche assay that measures total antibodies against the nucleocapsid protein of SARS-CoV-2. Given that our study focused on quantitative relationships between antibody levels and neutralization efficacies, we excluded from the analyses all samples that were negative for SARS-CoV-2 antibodies in at least 5 of the 8 immunoassays. Our final test set was composed of 85 samples (57 serum and 28 plasma). The Ortho test was only performed with the 57 serum samples due to assay incompatibility with plasma.

**Figure 1.**
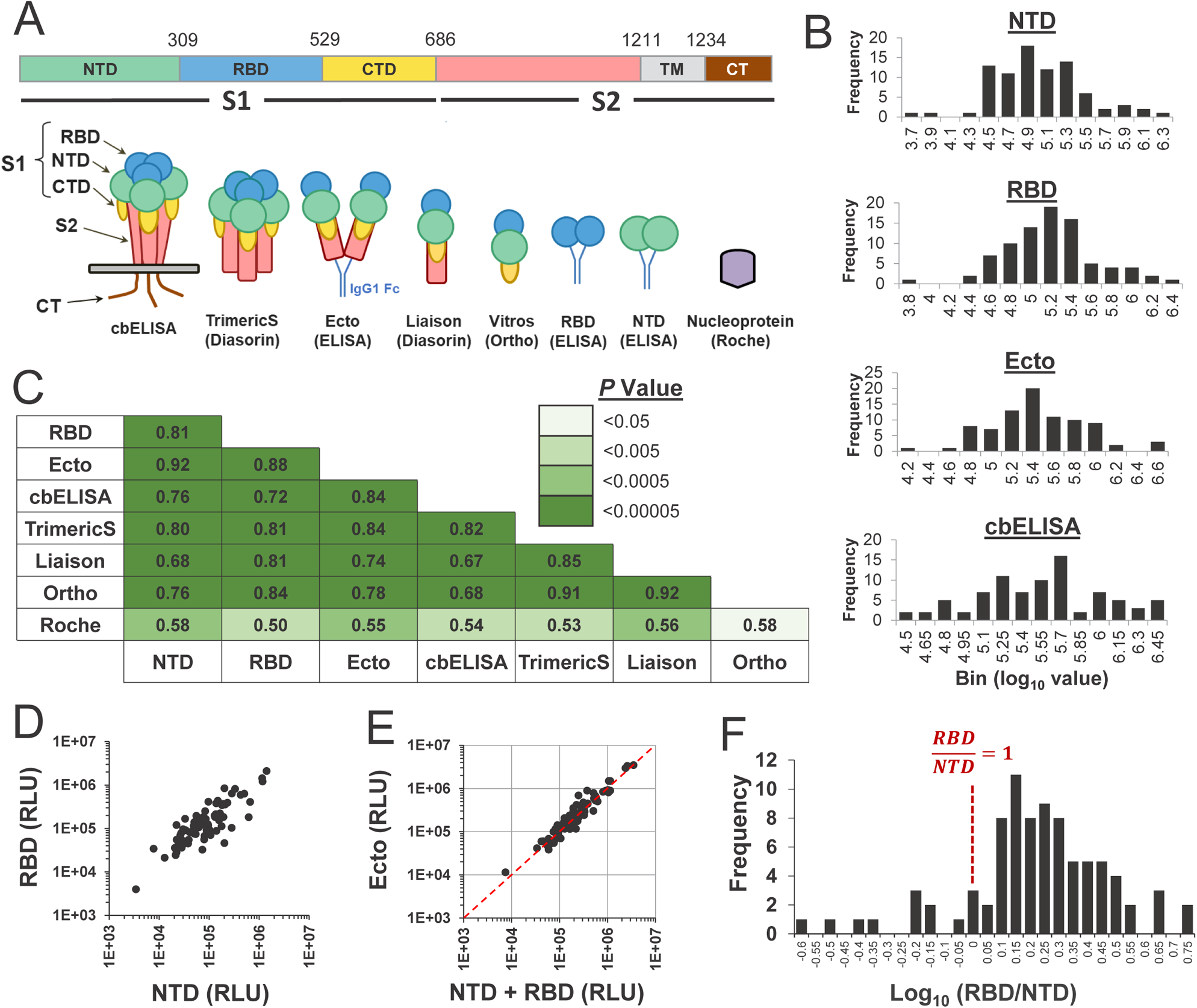
Comparison of immunoassays that apply different components of the SARS-CoV-2 spike protein as capture antigens. **(A)** Top, domains of the spike protein. NTD, N-terminal domain; RBD, receptor-binding domain; CTD, C-terminal domain; TM, transmembrane domain; CT, cytoplasmic tail. Bottom, schematic of the constructs used as capture antigens in this study. (B) SARS-CoV-2 antibody levels in 85 convalescent serum and plasma samples were tested using the RBD, NTD and Ecto ELISAs, as well as the cell-based ELISA. Distributions of the log_10_-transformed values are shown (see similar plots for commercial assays in **Supp. Fig. S1C**). **(C)** Spearman rank correlation coefficients between values measured in the eight immunoassays. Cells are colored by the *P* values determined in a two-tailed test. **(D)** Correlation between values in ELISAs that apply the NTD and RBD as capture antigens. RLU, relative light units. **(E)** Correlation between the sum of the values measured in the ELISAs that apply RBD and NTD, and values measured in the ELISA that applies Ecto as the capture antigen. The dashed red line describes a 1:1 relationship. **(F)** Distribution among the 85 convalescent samples of the ratio between log-transformed values measured in the RBD and NTD assays.

**Table 1.** Immunoassays and capture antigens used in this study.

| **Assay** | **Samples tested**<br>**(serum, plasma)** | **Capture antigen**<br>**used in assay** | **Assay type** | **Ig isotype**<br>**detected<sup>a</sup>** |
| --- | --- | --- | --- | --- |
| **NTD**<br>**(ELISA)** | 85 (57, 28) | NTD | In-house ELISA | IgG, IgM, IgA |
| **RBD**<br>**(ELISA)** | 85 (57, 28) | RBD | In-house ELISA | IgG, IgM, IgA |
| **S1/S2**<br>**(ELISA)** | 85 (57, 28) | S1/S2<br>ectodomain | In-house ELISA | IgG, IgM, IgA |
| **Cell-based**<br>**ELISA** | 85 (57, 28) | Full-length S1/S2<br>trimers | In-house ELISA | IgG, IgM, IgA |
| **TrimericS**<br>**(DiaSorin)** | 85 (57, 28) | Trimeric S1/S2<br>ectodomain | Commercial<br>assay | IgG |
| **Liaison**<br>**(DiaSorin)** | 85 (57, 28) | S1/S2<br>ectodomain | Commercial<br>assay | IgG |
| **Vitros**<br>**(Ortho)** | 60 (57, -) | S1 | Commercial<br>assay | IgG |
| **Roche** | 71 (43, 28) | Nucleocapsid | Commercial<br>assay | IgG, IgM, IgA |

<sup>a</sup> The secondary antibody used in the NTD, RBD and Ecto ELISAs and the cell-based ELISA targets the human kappa light chain.

The RBD, NTD and Ecto ELISAs, as well as cbELISA showed a normal distribution of their log_10_-transformed values (see **Fig. 1B** and results of a Shapiro-Wilk test in **Supp. Fig. S1B**). The log_10_-transformed values of the Liaison and TrimericS tests were also normally distributed, whereas the Roche test showed no evidence for normality (**Supp. Fig. S1B** and **S1C**). We compared values measured in the different assays using the non-parametric Spearman rank test. Strong correlations were observed between values measured in the assays that apply different spike components as the capture antigens, whereas correlations with the nucleocapsid-based Roche assay were less strong (**Fig. 1C** and **Supp. Fig. S2**). Interestingly, a strong association was observed between the content of antibodies against the non-overlapping NTD and RBD of spike (**Fig. 1D**).

Previous studies have suggested that the majority of spike-targeting antibodies elicited after infection or vaccination target the RBD and NTD, whereas antibodies that target the CTD and S2 are less common (25, 26). We used our in-house ELISA assays to determine the relative amounts of antibodies against the RBD and NTD of spike. Since equimolar concentrations of the NTD, RBD and Ecto proteins were used for capture in our ELISAs, we first compared for each sample the sum of the ELISA values measured in the RBD and NTD assays with the value measured for Ecto. We observed that for each sample, the sum of the NTD and RBD values was comparable to that of Ecto, indicating that antibodies against the NTD and RBD account for the vast majority of all S1/S2-targeting antibodies (**Fig. 1E**). We further compared the level of antibodies that target the RBD and NTD by calculating for each patient the ratio between the values in these assays. Greater binding activity to the RBD than NTD was observed, with a mean RBD-to-NTD ratio of 1.8 and standard deviation of 0.99 (**Fig. 1F**). Thus, the ratio of RBD-to-NTD antibodies was relatively constant in different subjects, ranging between 1 and 3 in 78% of cases. Only 2% of the samples showed two-fold or higher binding to the NTD, and only 8% of the samples showed more than 3-fold higher binding to the RBD. The proportion of RBD- or NTD-targeting antibodies (relative to all spike-targeting antibodies) did not vary with the interval between infection and the time of sample collection (**Supp. Fig. S3**).

Therefore, the levels of antibodies elicited against the RBD and NTD are highly correlated. Antibodies targeting the two domains exhibit a relatively constant relationship and account for the vast majority of all anti-spike antibodies elicited.

### Different domains, subunits and oligomeric forms of spike show similar abilities to estimate the neutralization efficacy of COVID-19 convalescent samples

Neutralizing antibodies mainly target the RBD and NTD of spike (9, 16, 17). Previous studies have shown that the levels of antibodies against different forms of spike (including S1, S1/S2 and RBD) correlate well with the neutralization capacity of the samples (27–31). However, the relationship between the neutralization activity of the polyclonal response in each individual and its domain specificity, as well as the variation in this relationship between different hosts are still poorly characterized. To address these questions, we compared the neutralization efficacy of the convalescent samples and their binding to the capture antigens that represent different domains and forms of spike. To quantify neutralization, we first used a replication-defective pseudovirus that contains the spike protein of SARS-CoV-2. For this purpose, we generated vesicular stomatitis virus (VSV) pseudovirions that encode the firefly luciferase gene in place of the native VSV-G glycoprotein gene and are pseudotyped with SARS-CoV-2 spike (VSV-SARS2-S) (5, 32). Residual infectivity of the pseudovirus in the presence of sera was measured using Vero-E6 target cells. The calculated dilution of sera at which virus infectivity was reduced two-fold is reported as the IC_50_ value. The log_10_-transformed IC_50_ values were consistent with a normal distribution (*P* value = 0.329 in a Shapiro-Wilk test), with a median IC_50_ corresponding to a dilution of 1:914 (**Fig. 2A**). Immunoassay values from each of the 8 tests were compared with the measured IC_50_ values (**Fig. 2B**). As expected, strong correlations were observed for all spike-based assays, as determined by the Spearman rank correlation coefficient, with *P* values lower than 0.000002 for all assays other than the nucleocapsid-based Roche test (**Fig. 2C**). These findings correspond with previous studies, which showed that spike-based immunoassays exhibit better correlations with IC_50_ values than the nucleocapsid-based Roche assay (33–35).

**Figure 2.**
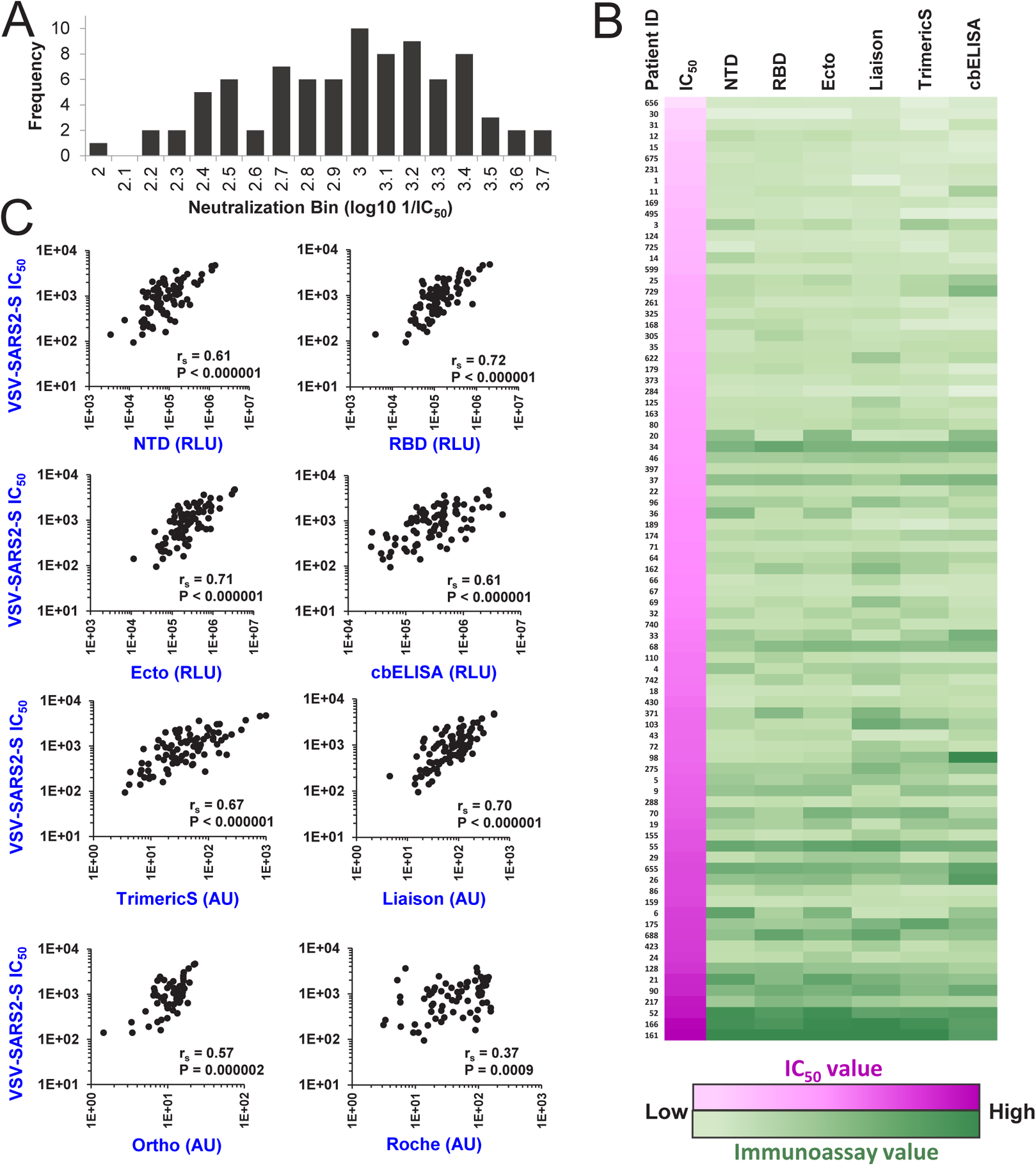
Relationships between immunoassay values of COVID-19 convalescent samples and their neutralization of spike-containing pseudovirus. **(A)** Neutralization titers of the serum or plasma samples were measured using replication-defective pseudovirus that contains the spike protein of SARS-CoV-2. Data describe the distribution of the log_10_-transformed IC_50_ values. **(B)** Comparison of neutralization and immunoassay values. All 85 samples are ordered by their neutralization titers (color-coded in shades of pink, with low values in lighter shades). Values measured in immunoassays are color-coded in shades of green. **(C)** Correlations between immunoassay values and neutralization titers. r_S_, Spearman correlation coefficient. *P* value, two-tailed test.

While there is a clear relationship between immunoassay values and neutralization efficacies in all comparisons, our primary question concerned the relative balance of these two factors in each assay, requiring a different metric. To better assess the relative abilities of the different capture antigens to estimate neutralization efficacies of the donor samples we used the precision metric. Precision was calculated by the ratio between the number of samples that are positive for neutralization at the selected threshold and the number of samples that are positive for SARS-CoV-2 antibodies by their immunoassay values. The level of precision was determined for subsets of the samples with progressively higher immunoassay thresholds for inclusion; from the 0^th^ percentile (all samples are included in the test) to the 98^th^ percentile (only samples with the top 2% of immunoassay values are included). Specific thresholds for neutralization were tested first, whereby a sample was considered neutralization-positive if the IC_50_ value was greater than a dilution of 1:500 or 1:2,000 (**Fig. 3A**). As expected, the use of samples from higher immunoassay percentiles resulted in higher precision. Differences between the immunoassays were more pronounced when the high neutralization threshold (1:2,000) was set. At this threshold, precision of the nucleocapsid-based Roche assay was low. Surprisingly, the cbELISA, which measures binding of antibodies to the native membrane-associated form of spike, also exhibited lower precision than other spike-based assays.

**Figure 3.**
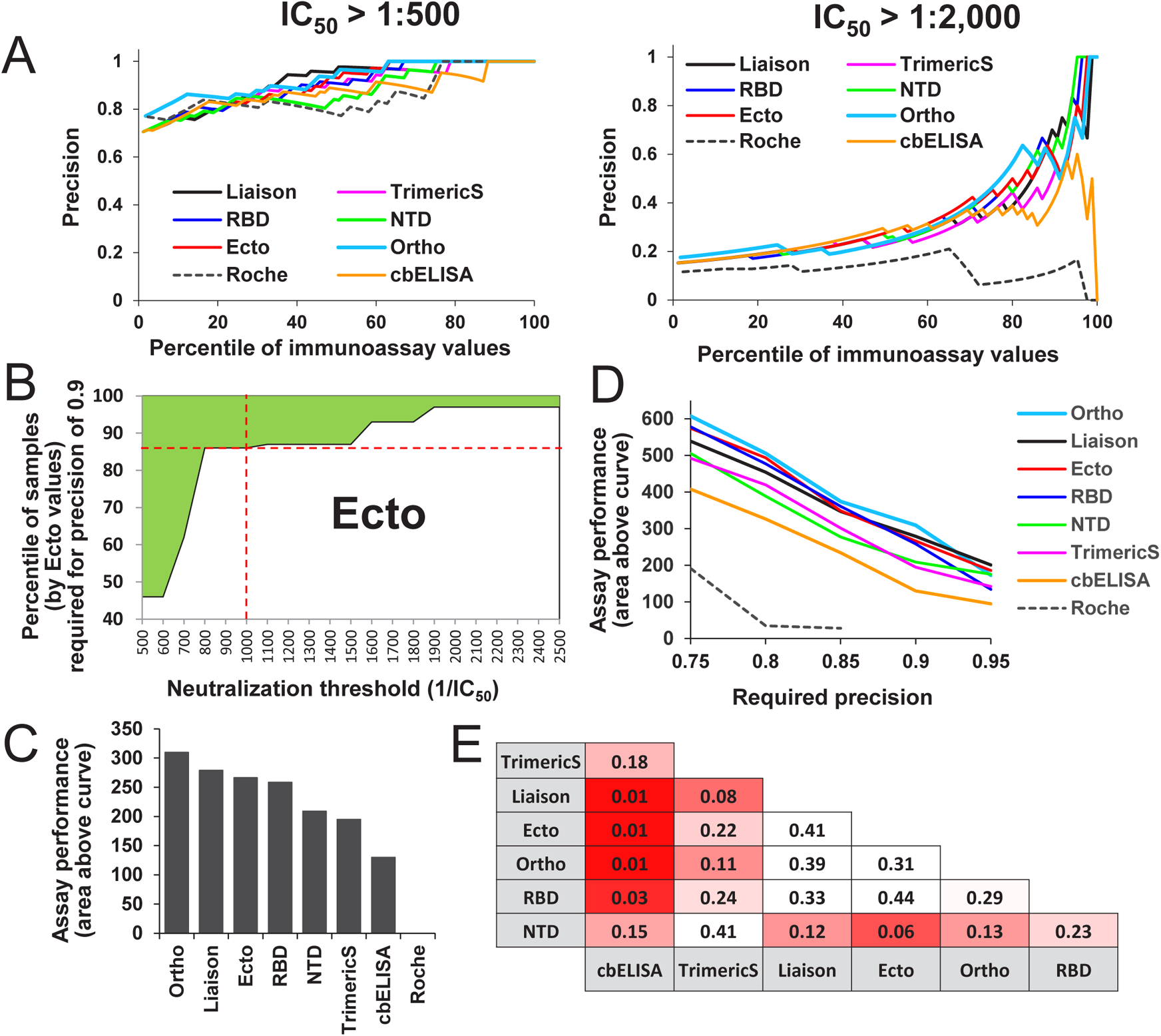
Precision of immunoassays to estimate the neutralization efficacy of COVID-19 convalescent serum and plasma. **(A)** Calculations of precision across different immunoassay percentiles. Precision was calculated as the number of samples with an IC_50_ greater than the defined threshold relative to the number of samples in the immunoassay percentile tested. Each data point describes precision of an immunoassay to predict neutralization at the indicated IC_50_ threshold using the indicated percentile of samples. **(B)** The area above the curve metric. The border between the shaded and unshaded areas describes the percentiles of Ecto values required to predict neutralization at the indicated thresholds with a precision of 0.9. For example, the intersection between the red lines indicates that, to predict with a precision of 0.9 for a threshold IC_50_ of 1:1,000, samples with Ecto values in the 85^th^ percentile should be used. The shaded area describes all neutralization threshold-percentile combinations that yield a precision of 0.9 or higher. **(C)** The area above the curve shown in panel B calculated for all immunoassays, based on a required precision of 0.9. **(D)** Calculations of the area above the curve for required precision levels of 0.75 to 0.95. **(E)** Statistical significance of the differences between predictive capacity of the immunoassays. The area above the curve was calculated for all immunoassays for a precision of 0.9. Significance of the difference between predictive capacity of any two assays was determined by a permutation test. *P* values of the one-sided test are shown. Cells are color coded by the *P* values calculated.

Given that performance of the assays can vary at each neutralization threshold, we sought to generate a metric that would describe performance across a range of neutralization thresholds. To this end, we first calculated for each threshold (from 1:500 to 1:2,500) the minimal immunoassay percentile required to achieve a precision level of 0.9 (see boundary line for Ecto ELISA as example in **Fig. 3B** and all assays in **Supp. Fig. S4**). The area above the boundary line indicates the percentile-threshold combinations that yield a precision of 0.9 or higher, which allows us to compare overall performance characteristics between immunoassays; the greater the area, the higher the ability of the assay to predict neutralization across all IC_50_ thresholds. The highest performance was observed for the Ortho, Liaison, Ecto and RBD assays, followed by NTD, TrimericS and cbELISA (**Fig. 3C**). Since the Roche assay did not achieve a precision of 0.9, the areas above the curve could not be computed. We then calculated the area above the curve when the required precision was set at levels ranging between 0.75 and 0.95. For most precision requirement levels in this range, the lowest performance was observed for the Roche assay, followed by cELISA, with modestly better performance for the TrimericS and NTD (**Fig. 3D**). All other assays exhibited similar performance across the different precision requirements. To determine statistical significance of the differences between performance of any two assays, we performed a permutation-based test (see Materials and Methods section). Briefly, for each pair of assays compared, we measured the area above the curve and calculated the difference. We then permuted for each patient sample the immunoassay identifiers, the area above the curve was recalculated for both immunoassays and the difference determined. The fraction of the times the difference was greater using the permuted values relative to the unpermuted values was calculated as the *P* value. Significant differences for a one-sided test (*P* values lower than 0.05) were observed between the cbELISA and all other spike-based assays. The NTD and TrimericS assays showed moderate differences from other assays; however, they were not significant at the 95% confidence level (**Fig. 3E**). Therefore, the ability of cbELISA (i.e., the full-length membrane-bound form of spike) to predict neutralization was significantly lower than that of all other assays that apply isolated domains of the protein as capture antigens.

To independently validate the above findings, we also measured neutralization titers for 24 of the serum samples using infectious SARS-CoV-2 under BSL-3 conditions, and correlated those findings with immunoassays values. Virus-induced cytopathology was used to detect infection. The dilution of serum at which cytopathic effects were observed in fewer than 50% of the wells was determined, and data were fit to a regression model to calculate the precise IC_50_ value. For three of the samples, the IC_50_ was not achieved at the lowest dilution of the serum used (1:40); the remainder showed a range of IC_50_ values, with a median dilution of 1:212 (**Fig. 4A**). A strong correlation was observed between the neutralization titers of the sera measured using the replicative SARS-CoV-2 and the VSV-based pseudovirus that contains the spike protein (**Fig. 4B**). As expected, IC_50_ values in the pseudovirus assay were higher than those measured using infectious virus, since the former measures the dilution at which 50% of virus infectivity is reduced whereas the latter assay measures the dilution at which more than 50% of wells show complete neutralization of all input virus.

**Figure 4.**
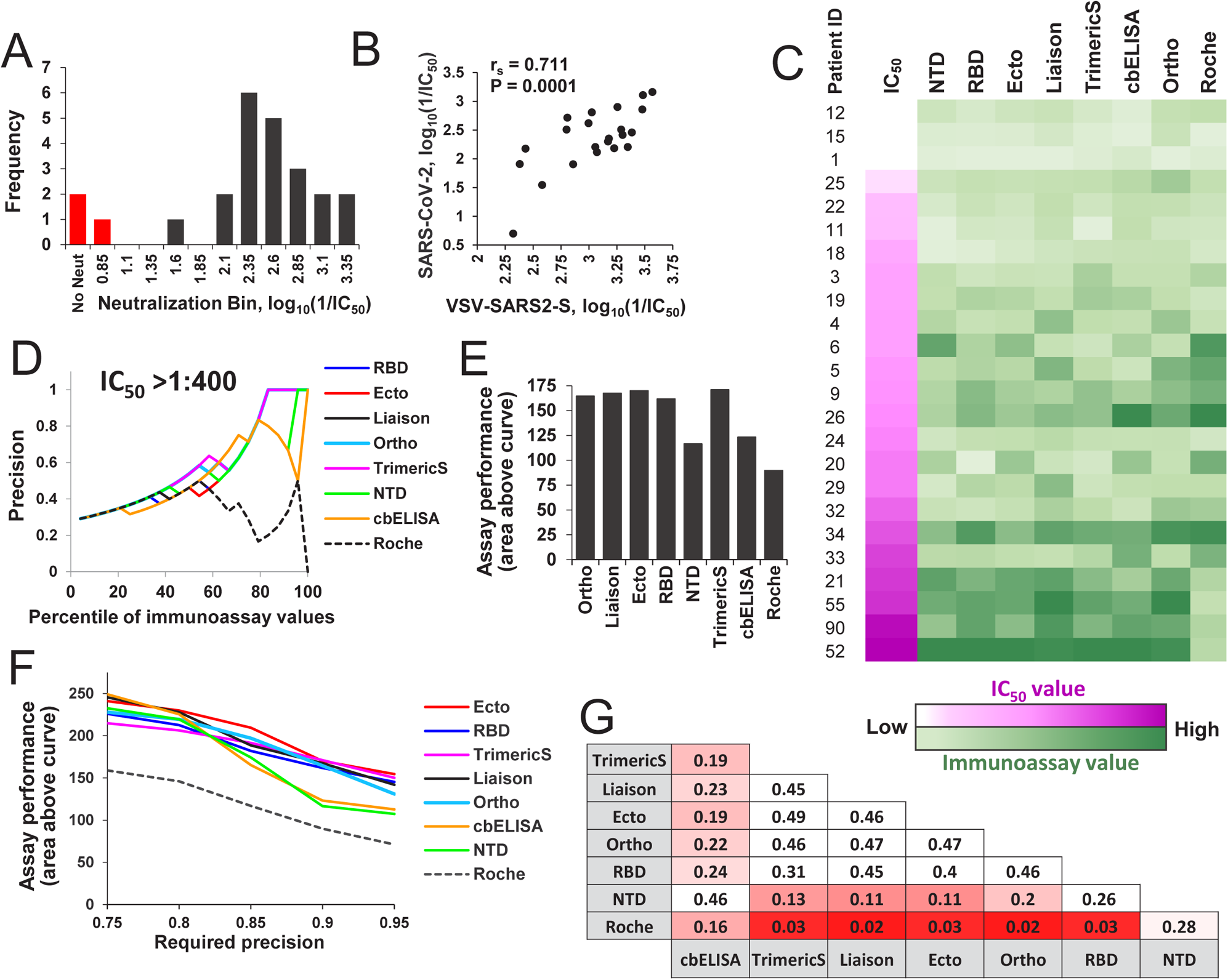
Immunoassay-based estimations of replicative SARS-CoV-2 neutralization. **(A)** Twenty-four serum samples were tested for their neutralization of replicative SARS-CoV-2. The distribution of IC_50_ values is shown. For three samples, the IC_50_ was not achieved at the lowest dilution (1:40) of the serum (columns colored in red). **(B)** Correlation between IC_50_ values of the 24 serum samples, as measured using replication-competent SARS-CoV-2 and the VSV-SARS-S pseudovirus. **(C)** Comparison of neutralization and immunoassay values. Samples are ordered by their neutralization efficacy of replicative SARS-CoV-2 (color-coded in shades of pink, with low values in lighter shades). Values measured in immunoassays are color-coded in shades of green. **(D)** Precision of immunoassays to estimate SARS-CoV-2 neutralization at an IC_50_ threshold of 1:400 using different percentiles of the samples based on their immunoassay values. **(E)** Area above the curve calculated for a required precision of 0.9. **(F)** Performance of immunoassays to estimate neutralization at precision levels of 0.75 to 0.95. **(G)** Statistical significance of the differences between predictive capacities of the immunoassays, as determined by a permutation test. Calculations apply the area above the curve computed for a required precision of 0.9. *P* values of the one-sided test are shown. Cells are color coded by the *P* values.

We compared immunoassay values of the samples with their neutralization efficacies of replicative SARS-CoV-2 (**Fig. 4C**). Strong correlations were observed for all spike-based assays (**Supp. Fig. S5**). Precision analyses using an IC_50_ threshold of 1:400 demonstrated considerable differences between performance of the assays (**Fig. 4D**). Comparison of the overall performance of the immunoassays across neutralization thresholds of 1:50 to 1:500 (using the area above the curve metric with a required precision of 0.9) showed a similar pattern to the pseudovirus-based measurements (compare **Fig. 4E** and **Fig. 3C**); the poorest performance was observed for the Roche assay, followed by NTD and cbELISA. All other assays performed similarly well. Comparison of assay performance at precision levels of 0.75-0.95 showed modest differences between cbELISA or NTD and all other spike-based assays (**Fig. 4F**); however, these differences did not reach a significance level of 0.05 (**Fig. 4G**)

Taken together, these results demonstrate that performance of immunoassays based on RBD, S1, or monomeric and dimeric forms of S1/S2 to estimate the neutralization efficacy of each sample was similar. Modestly lower predictive capacities are observed when NTD and the full-length form of spike (as measured by cbELISA) are used as the capture antigens. Further, comparison of the precision of the immunoassays to predict neutralization using pseudovirions containing SARS-CoV-2 spike or replication-competent viruses yield roughly similar findings.

### COVID-19 convalescent samples exhibit a similar level of relative neutralization potency

The above results show that different forms of spike used as capture antigens (NTD, RBD, S1 or Ecto) can estimate neutralization with similar precision. Furthermore, the relationship between the levels of NTD and RBD antibodies is relatively conserved in different individuals; these antibodies compose the vast majority of the antibodies generated against spike. We asked whether the neutralization efficacy increases with higher proportions of RBD-or NTD-targeting antibodies (relative to all spike-targeting antibodies). Comparison of the RBD-to-Ecto or NTD-to-Ecto ratios with the neutralization efficacy of the samples showed no evidence for a relationship between these variables (**Fig. 5A** and **5B**). Similarly, the RBD-to-NTD ratio was not associated with the neutralization efficiency of the samples (**Fig. 5C**). These findings indicate that convalescent samples with high neutralizing activity do not contain a higher proportion of antibodies that target the RBD or NTD.

**Figure 5.**
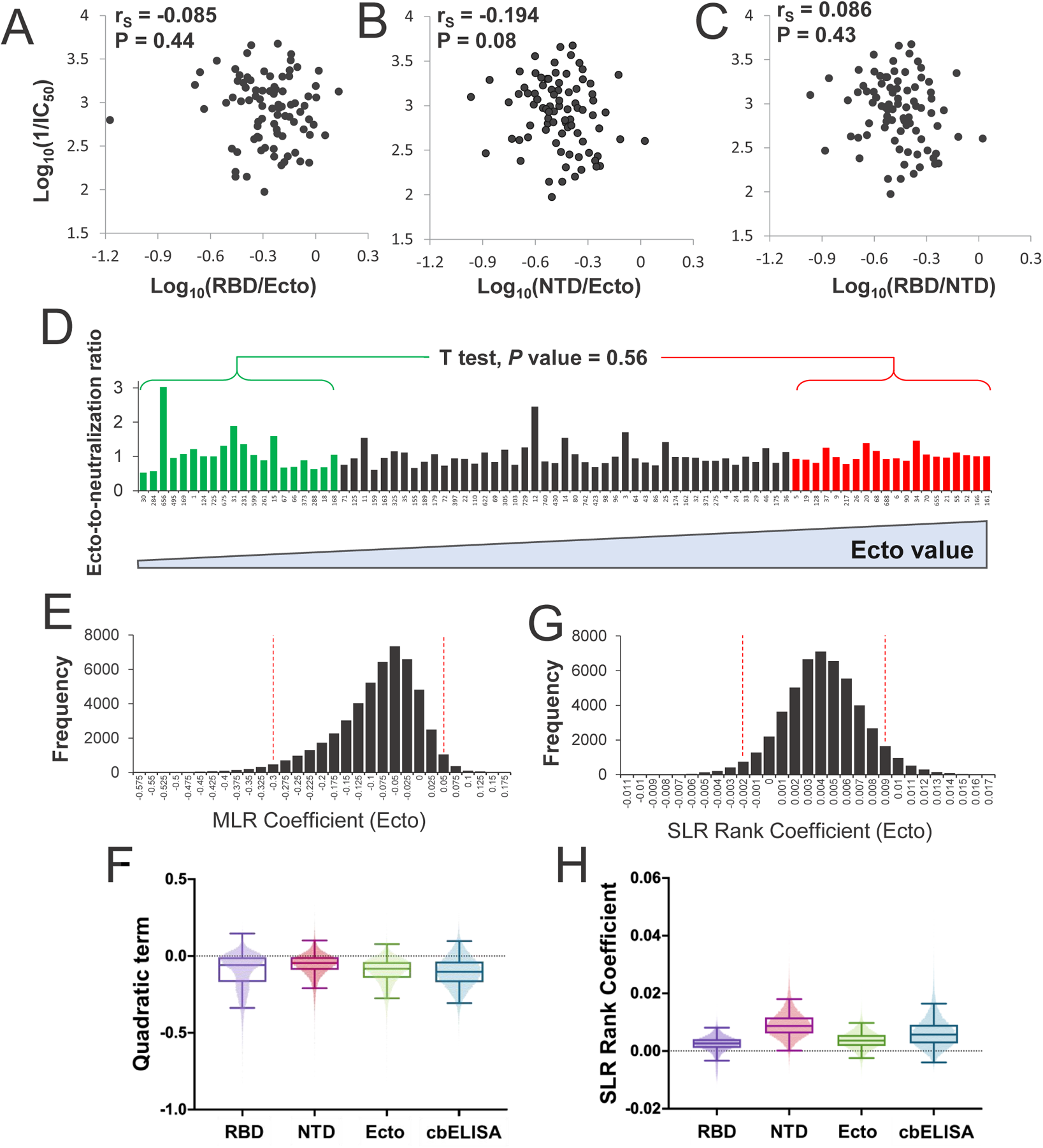
Relationship between the relative neutralizing potency of convalescent samples and their content of spike-specific antibodies. **(A,B)** Comparison of the relative RBD or NTD binding values (expressed as a fraction of the Ecto value) and neutralization efficacy of the samples. (C) Comparison of the ratio between RBD and NTD values and neutralization efficacy. **(D)** IC_50_ values of convalescent samples and their immunoassay values were log_10_-transformed and adjusted to a scale of 0.1 to 1. The ratio between the Ecto ELISA value and the IC_50_ of each sample was calculated for all samples. Patient samples are arranged by increasing Ecto values, from left to right. The ratios calculated for the 20 samples with the lowest and the 20 samples with the highest Ecto values were compared using an unpaired T test; the *P* value for a two-tailed test is indicated. **(E)** Bootstrap distribution for quadratic term in MLR describing the relationship between log-Ecto values and log-IC_50_ values, evaluating evidence for a non-constant ratio relationship. A 95% bootsrap confidence interval was determined from bootstrap sample quantiles. **(F)** The quadratic term was calculated by 50,000 iterations of bootstrap resampling for NTD, RBD, Ecto and cbELISA data. The boxed area shows the second and third quartiles. Whiskers describe the range for two standard deviations. **(G)** Bootstrapped rank regression. The rank order of Ecto values for all 85 samples was determined as well as the ratio between the ranks of the Ecto value and IC_50_. A simple linear regression model was fit to the relationship between the two variables. A bootstrapping procedure was performed to estimate the slope coefficient. The bootstrap distribution and corresponding 95% confidence interval are shown. **(H)** The rank regression coefficient was calculated by bootstrap resampling using NTD, RBD, Ecto or cbELISA data.

A large proportion of spike-targeting antibodies elicited by infection are non-neutralizing (36, 37). A recent study has shown that infected and immunized hosts with high levels of spike-specific antibodies generate a significantly higher proportion of non-neutralizing antibodies than individuals with lower levels of anti-spike antibodies (38). To explore this relationship in our samples, we implemented a model to examine evidence for a variable ratio between immunoassay values and neutralization efficacy. Two computational approaches were used; the first looks for non-log-linearity in the relationship between neutralization and immunoassay tests, whereas the second considers their rank-ratios and examines evidence for a systematic change over the ranks of the immunoassay results.

To compare the variables and avoid a bias related to the dynamic ranges of the values, we corrected the log_10_-transformed immunoassay and neutralization IC_50_ values to the same scale by adjustment to a range from 0.1 to 1. For each sample we calculated the ratio between the immunoassay value and the IC_50_ value (see analysis of the Ecto ELISA data in **Fig. 5D**). This ratio was compared between the 20 samples with the lowest immunoassay values and the 20 samples with the highest immunoassay values. Evaluation of these results did not find significantly different ratios in the two groups (see *P* value for an unpaired T test in **Fig. 5D**). A similar lack of a significant difference was observed when the RBD and NTD were used as capture antigens (**Supp. Fig. S6**). However, the cbELISA results suggested a higher ratio (i.e., a lower relative neutralization efficacy) for the samples with high antibody levels.

To further explore whether the immunoassay-to-neutralization ratio shows any indication of dependence on the immunoassay value, we examined the variability in this ratio by looking for non-linearity in their log-relationship using all 85 samples. The null hypothesis tested was that the log-scale relationship between these variables should be linear, which was tested by considering a quadratic term for immunoassay results in a multiple linear regression (MLR) model. While the data appeared well modelled directly on a log-10 scale, to eliminate concerns about distributional assumptions, the regression coefficient was bootstrapped, and the corresponding 95% confidence interval determined. We first analyzed the results of the Ecto assay. As shown in **Fig. 5E**, an MLR slope value of 0 (i.e., lack of a quadratic effect, leaving a linear increase in neutralization activity for a given increase in binding) lies within the 95% confidence interval, so we fail to reject the null hypothesis that the variables follow a ratio relationship. Similar analyses of the data from the NTD, RBD and cbELISA tests also failed to show evidence at the 95% level to support a non-linear relationship between immunoassay values and neutralization (**Fig. 5F**).

We also applied a rank-based approach, whereby immunoassay and neutralization values were transformed to their ranks (from 1 to 85). A simple linear regression (SLR) coefficient was then fitted to the relationship between the immunoassay rank value and immunoassay-to-neutralization rank-ratio, and bootstrapping was applied once more to produce 95% confidence intervals. The null hypothesis tested was that a slope of zero exists for this relationship. Again, no evidence was observed to support the notion that the ratio between Ecto values and neutralization varies across different levels of S1/S2-targeting antibodies (**Fig. 5G**). A similar bootstrapping analysis of the rank values for the RBD, NTD and cbELISA failed to demonstrate a non-zero slope that would indicate a linear relationship between the two variables (**Fig. 5H**).

Given the sample size (n=85), the presence of a strong relationship between neutralization fraction and antibody binding activity seems unlikely. Nevertheless, we do observe negative non-significant coefficients for the quadratic effect of log-binding activity on neutralization levels, and positive non-significant coefficients of for the linear relationship between binding activity and the rank ratio of binding to neutralization (**Fig. 5, G** and **H**). Both of these results indicate the plausibility of a weak relationship between the neutralization ratio and binding activity measures, in which higher binding activity could be associated with lower proportional neutralization activity, but the magnitude of such an effect is likely to be limited.

## DISCUSSION

Over the course of the COVID-19 pandemic, our understanding of the antibody response against SARS-CoV-2 has evolved. Initial investigations suggested that most neutralizing antibodies elicited by infection or vaccination target the RBD (9, 39). More recent studies have shown a co-dominance of antibodies that target the RBD and NTD (25, 26). Proteomic deconvolution studies of the IgG repertoire in COVID-19 convalescent patients suggested that the bulk of the neutralizing response targets epitopes outside the RBD (40). To better understand the target specificity of the response in different individuals, we analyzed the relative level of antibodies against different domains, subunits and oligomeric forms of spike in COVID-19 convalescent samples. Our findings suggest the model shown in **Fig. 6**. A polyclonal antibody response is elicited in each infected individual against multiple domains of spike. High variation is observed between individuals in the amounts of antibodies generated; however, there is limited variation in the proportion of antibodies against the RBD and NTD (relative to all anti-spike antibodies). Similarly, limited variation is observed in the relationship between the amounts of antibodies against the RBD and NTD, with a ratio ranging between 1 and 3 in 78% of subjects. Importantly, the relative potency of the response (i.e., the level of neutralizing activity relative to the level of antibodies generated) is also constant in different individuals. Thus, the domain specificity and relative inhibitory activity of the response is conserved among individuals, with the main variation being the total amount of the antibodies produced.

**Figure 6.**
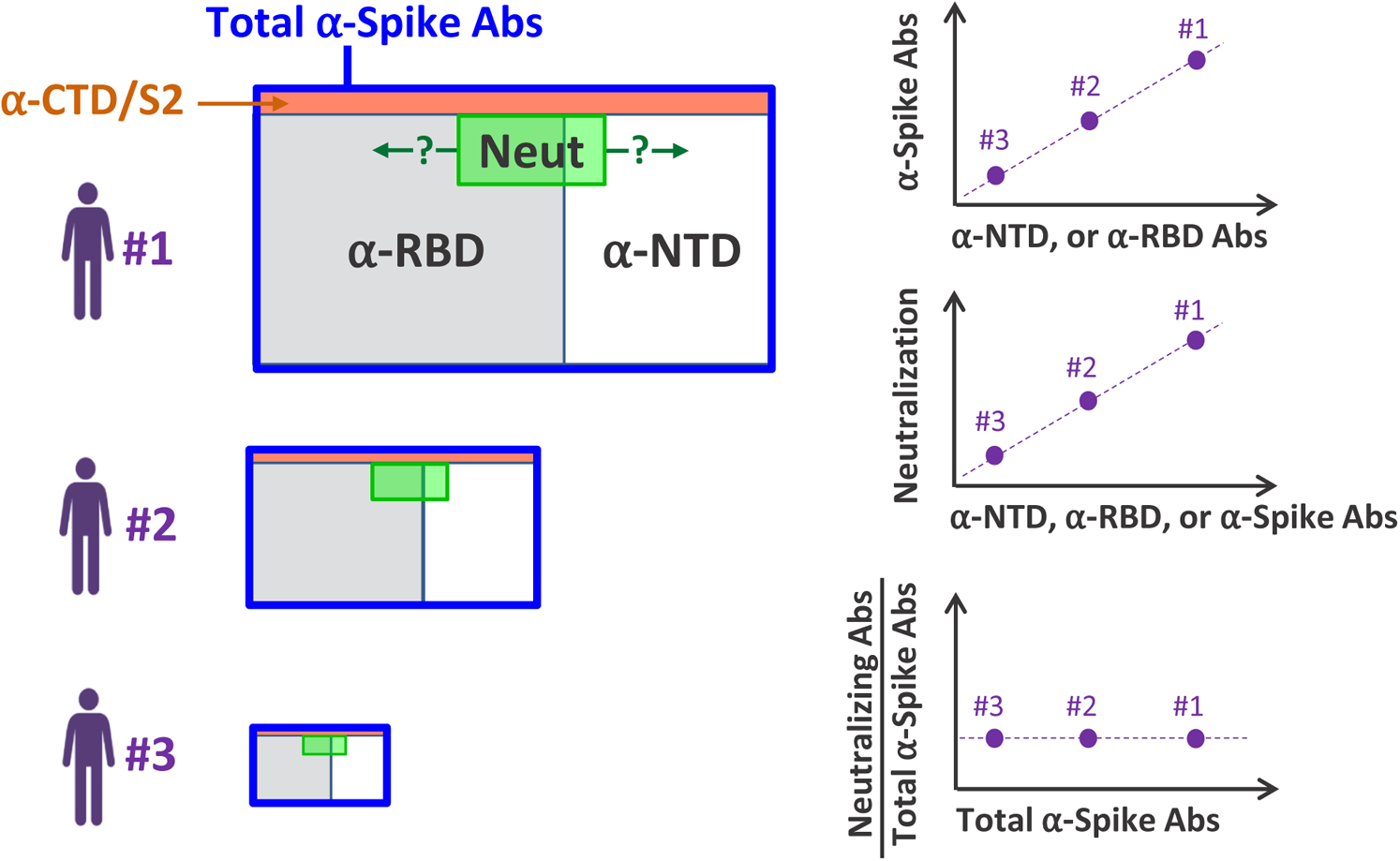
Model of the polyclonal antibody response elicited against the SARS-CoV-2 spike protein. SARS-CoV-2-infected individuals generate different amounts of spike-targeting antibodies (represented by the size of the blue rectangles). The fraction of antibodies that target the RBD or NTD is constant in different individuals, with a mean RBD-to-NTD ratio of 1.8. The relative neutralization potency (represented by the green-shaded area) is also similar in different individuals. Question marks indicate the yet unknown domain distribution of the fraction of antibodies that contains neutralizing activity (i.e., positioning of the green-shaded area).

Multiple commercial immunoassays have been developed that apply different components of the spike protein as the capture antigen (35, 41–44). Spike-based assays have been shown to estimate well the neutralization efficacy of convalescent serum and plasma samples (27–30). They are thus applied as qualitative measures of immunization status and can potentially serve as indirect measures of the efficacy of the anti-SARS-CoV-2 humoral response. Our studies suggest that any soluble form of spike that contains the RBD or NTD can serve as a capture antigen to accurately determine the immunization status of the individual and potentially the efficacy of the anti-SARS-CoV-2 humoral response. The constant proportion of antibodies against different spike domains also explains the ability of immunoassays that use various forms of the protein as capture antigens to predict neutralization. Indeed, our findings suggest that determinations of neutralizing titers based on serological tests do not require native forms of the protein as the capture antigen; RBD exhibits a similar predictive capacity to that of S1 or S1/S2, with only modestly lower performance for NTD. Thus, inclusion of S2 or trimerization of the protein to mimic the native form of spike does not improve the ability to estimate the amount of neutralizing antibodies. In fact, the poorest performance was observed for the full-length, membrane-bound form of the protein measured by cell-based ELISA. The lower predictive capacity of the cbELISA may result from detection of non-neutralizing antibodies that may recognize the native form of spike (37). Alternatively, differential post-translational processing of spike in the HOS cells (relative to the human embryonic kidney 293T cells used to produce the recombinant proteins for these assays) may affect antigenicity of this protein (45).

We were surprised to discover that subjects with different amounts of spike-specific antibodies contained a constant level of relative potency. Such results contrast with a recent study by Amanat et al., which suggested that convalescent samples that contain high amounts of spike antibodies (as measured by the Mount Sinai Laboratory COVID-19 ELISA IgG Antibody Test) contain a higher proportion of non-neutralizing antibodies that target the full-length ectodomain of spike (38). It should be noted that in their calculations, the authors analyzed the immunoassay-to-IC_50_ ratios using the raw values obtained in these tests. Unfortunately, such an approach can introduce a bias if the dynamic ranges of the two variables differ, which may impact the results of the analysis. To address this potential bias, we performed our calculations using ranks and values that were corrected to the same scale. Both approaches showed similar results, whereby the relative potency is constant in different samples, regardless of the amount of anti-spike antibodies generated. Future studies will reveal whether the target specificity of antibodies with neutralizing activity is also constant in different individuals and independent of the robustness of the response. Such studies are of particular importance in vaccinated individuals, to accurately quantify and characterize specificity of the antibody fractions that can protect from infection.

## MATERIALS AND METHODS

### Collection of plasma and serum from donors and patients

All blood donors were screened following the FDA guidance instructions under an institutional review board approved protocol (IRB #202003554). The consent signed by all donors allowed the use of blood samples for research purposes. Donors were identified and screened following FDA guidelines at the time they enrolled. Two study groups were assessed. The first is composed of 57 convalescent serum samples from subjects that had either been confirmed by reverse transcription polymerase chain reaction (RT-PCR) to be SARS-CoV-2 positive from a nasopharyngeal swab (n=51) or had signs or symptoms of COVID-19 and were found to be positive by serological testing (n=6). All donors except one had relatively mild COVID-19 symptoms; this donor was hospitalized for one day due to palpitations. Donor screening was performed at least 10 days after resolution of symptoms. At the time of plasma collection, serum samples were collected in serum separator tubes and allowed to clot for at least 30 minutes. Serum was then isolated, aliquoted and stored at −80°C until use. The second study group is composed of convalescent plasma collected from women hospitalized for delivery, who had previously been infected by SARS-CoV-2, as confirmed by a SARS-CoV-2-positive PCR (n=7) or positive serology test (n=21). Samples were collected in EDTA-containing tubes, aliquoted and frozen until use at −80°C.

### Cells lines

Vero-E6 cells, human embryonic kidney (HEK) 293T cells and human osteosarcoma (HOS) cells were obtained from the American Type Culture Collection (ATCC). Cells were cultured in Dulbecco’s modified Eagle medium (DMEM) supplemented with 2-10% fetal calf serum (FCS) and 1% penicillin/streptomycin. All cells were maintained in a humidified incubator at 37°C and 5% CO_2_.

### Recombinant proteins and their production

Capture antigens that contain different spike protein components were generated. The NTD, RBD or ectodomain of S1/S2 (Ecto) antigens were fused to the Fc region of human IgG1, rendering them dimeric. NTD and RBD contain amino acids 1-309 and 310-529 of spike, respectively. The Ecto protein contains the entire ectodomain of spike (amino acids 1-1274). To abrogate the furin cleavage site in Ecto, we substituted the Arg-Arg-Ala-Arg motif at position 683-686 with Ser-Ser-Ala-Ser. All proteins were produced by transient transfection of 293T cells using polyethyleneimine (PEI), as previously described (46). Proteins were harvested in 293S ProCDM and purified using Protein A beads. Eluted products were dialyzed against phosphate buffered saline (pH 7.4). All proteins were analyzed by SDS-PAGE and visualized by silver staining to verify their molecular weight and purity.

### ELISA using RBD, NTD and S1/S2 as capture antigens

The RBD, NTD and Ecto recombinant proteins were used as capture antigens in an enzyme-linked immunosorbent assay (ELISA). Briefly, proteins were suspended in PBS at a concentration of 25 nM (2 μg/mL of NTD, 1.37 μg/mL of RBD and 5 μg/mL of Ecto) and incubated overnight in protein-binding 96-well plates (PerkinElmer). The next day, wells were washed once with blocking buffer, composed of 140 mM NaCl, 1.8 mM CaCl_2_, 1 mM MgCl_2_, 25 mM Tris pH 7.5, 20 mg/ml BSA and 1.1% nonfat dry milk. Serum or plasma samples were diluted 1:500 (vol:vol) in blocking buffer, added to the wells and incubated for 45 min at room temperature. Samples were then washed four times with blocking buffer and a horseradish peroxidase (HRP)-conjugated secondary antibody that targets the kappa light chain of human IgG1 was added (diluted 1:1200 in blocking buffer). After incubation for one hour at room temperature, samples were washed 5 times with blocking buffer and 5 times with washing buffer (140 mM NaCl, 1.8 mM CaCl_2_, 1 mM MgCl_2_ and 25 mM Tris pH 7.5). HRP enzyme activity was measured by light emission using Supersignal West Pico Plus chemiluminescence detection reagents with a Synergy H1 microplate reader.

### Cell-based ELISA measurements of antibodies against SARS-CoV-2 spike

Binding of serum antibodies to SARS-CoV-2 spike expressed on HOS cells was measured using a previously-described cell-based ELISA system (22, 24). Briefly, HOS cells were seeded in white opaque 96-well plates (1.4 × 10^4^ cells per well) and transfected the next day with 80 ng per well of pCG1-SARS-2-S plasmid expressing SARS-CoV-2 spike using JetPrime transfection reagent. To quantify background binding of the antibodies to the cells, a similar number of wells was transfected using a negative control plasmid (ΔKS) that does not encode for a viral protein product (23). Three days after transfection, cells were washed with blocking buffer, and serum samples diluted 1,000-fold in blocking buffer were incubated with the cells for one hour at room temperature. Cells were then washed 5 times with blocking buffer and incubated for 45 minutes at room temperature with an HRP-conjugated goat anti-human kappa chain preparation that was diluted 1:1,200 in blocking buffer. Cells were then washed 5 times with blocking buffer and 5 times with washing buffer. HRP enzyme activity was measured by chemiluminescence with a Synergy H1 microplate reader.

### Commercial immunoassays to measure antibodies that target SARS-CoV-2 proteins

The DiaSorin Liaison SARS-CoV-2 S1/S2 IgG chemiluminescence assay detects IgG against spike subunits S1 and S2. Samples were analyzed according to the manufacturer’s guidelines on a DiaSorin Liaison XL automated chemiluminescence analyzer (DiaSorin, Saluggia, Italy). A signal of 15 arbitrary units (AU) per mL or higher is defined by the manufacturer as a positive result. The DiaSorin TrimericS IgG assay applies the soluble trimeric form of the S1/S2 subunits. Samples were analyzed on a DiaSorin Liaison XL automated chemiluminescence analyzer. A signal of 13 AU/mL is defined as a positive result. The Roche electrochemiluminescence immunoassay measures total immunoglobulins that target the SARS-CoV-2 nucleocapsid (N) protein. Samples were analyzed according to the manufacturer’s guidelines using the Elecsys Anti-SARS-CoV-2 assay on the Roche cobas e602 platform (Roche Diagnostics). A cut-off index (COI) of 1.0 or higher is defined by the manufacturer as a positive result. The Ortho COVID-19 IgG antibody test was performed on Ortho’s VITROS® system. The signal at cutoff (S/C) value as defined by the manufacturer is 1 unit or greater for a reactive sample. For simplicity, all values of the commercial immunoassays are reported in arbitrary units (AUs).

### Infection and neutralization of replication-competent SARS-CoV-2

SARS-CoV-2 strain USA-WA1_2020 was obtained from BEI Resources and maintained under biosafety level 3 (BSL-3) conditions. To propagate virus, Vero-E6 cells cultured in DMEM/FCS 2% were infected at a multiplicity of infection (MOI) of 0.001. Forty-eight hours after infection, supernatants were harvested, filtered through 0.45 μm pore-sized membranes, and frozen at −80°C until use. To quantify the amount of infectious virus, Vero-E6 cells were seeded in 96-well plates (1.5 · 10^4^ cells per well). The next day, serial dilutions of the virus were added to 8 replicate wells for each dilution and cytopathic effects were evaluated over the next 5 days. The median tissue culture infectious dose (TCID_50_) was used to quantify virus titer, which describes the dilution of the virus at which fewer than half of the replicate wells show cytopathic effects.

To measure neutralization, serial two-fold dilutions of the serum samples (ranging from 1:40 to 1:2,560) were prepared in DMEM/FCS 2%. Virus was added to the diluted serum at a final concentration of 25 TCID_50_ per well. Samples were incubated at room temperature for one hour and added to Vero-E6 cells seeded the day before in 96-well plates (1.5 × 10^4^ cells per well). Six replicate wells were used for each dilution. Cells were then cultured for 4 days at 37°C until infectivity was evaluated. The number of wells in which intact monolayers were present was assessed using an inverted light microscope. The 50% neutralizing titer (IC_50_) was calculated by the serum dilution at which 50% or more of the wells showed no cytopathic effects. To determine the precise IC_50_ value, the number of wells in which cytopathic effects were observed at each serum dilution was recorded. These values, along with the log-transformed dilution values were fit to a non-linear regression model using GraphPad Prism 8 to calculate the IC_50_ value.

### Production and neutralization of vesicular stomatitis virus pseudotyped by the SARS-CoV-2 spike protein

Vesicular stomatitis virus (VSV) pseudovirions bearing the SARS-CoV-2 spike protein (VSV-SARS2-S) were generated as previously described (5). Briefly, 293T cells were seeded in 100 mm plates (2.2 · 10^6^ cells per plate) and transfected 24 hours later by 16 µg of pCG1-SARS-2-S plasmid (a kind gift from Dr. Stefan Pöhlmann) using PEI transfection protocol (46). Twenty-four hours after transfection, cells were infected with a stock of VSV pseudovirus that encodes the firefly luciferase gene in place of the native VSV-G glycoprotein gene and contains the glycoprotein of Lassa virus (5). Six hours later, infected cultures were washed twice with phosphate buffered saline (PBS, pH 7.4) to remove input pseudovirions, and fresh DMEM/FCS 2% was added. Media was collected at 24-and 48-hours after infection, the supernatants were filtered through 0.45 μm pore-sized membranes and centrifuged at 5,380 × g for 16 hours at 4°C. The pellet was resuspended in PBS and centrifuged through a 20% sucrose cushion at 134,000 × g for 2 hours at 10°C. Pellets containing the pseudoviruses were resuspended in PBS and stored at −80°C until use.

For neutralization assays, two-fold serial dilutions of the serum samples were prepared in DMEM/5% FCS, ranging between 1:40 and 1:2,560. Viruses were added to the diluted serum at a concentration calculated to yield between 100,000 and 200,000 relative light units (RLUs) of luciferase activity per well. These values were determined to be within the linear range of virus input versus luciferase activity measured. Vero-E6 target cells were seeded the day before infection in 96-well white opaque flat-bottomed plates (1.5 × 10^4^ cells per well). The virus-serum or virus-plasma mixture was incubated for one hour at 37°C and added to the wells. Six replicate wells were used for each condition. Samples were then incubated for 24 h at 37°C, after which the media were removed and 35 µl of Passive Lysis buffer (Promega) was added to each well. Luciferase activity was recorded as a measure of viral infection, as previously described (24). Briefly, 100 µl of luciferin buffer containing 15 mM MgSO_4_, 15 mM KPO_4_ (pH 7.8), 1 mM ATP, and 1 mM dithiothreitol was added to each well, followed by 50 µl of 1 mM d-luciferin potassium salt (Syd Laboratories). Luminescence was detected using a Synergy H1 Hybrid reader (BioTek Instruments).

### Permutation test to compare precision of immunoassays

For each immunoassay, we obtained the curve that describes the required percentile of samples for each neutralization threshold to yield a precision of 0.9. The area above the curve was then determined, which describes all percentile-neutralization threshold combinations that yield a precision level higher than the minimum precision of interest (here, 0.9). This metric thus captures the precision of each assay across multiple neutralization thresholds. To test for significant difference between the area above the curve for any two immunoassays, we used a permutation test. The null and alternative hypotheses for a one-sided test can be stated as:

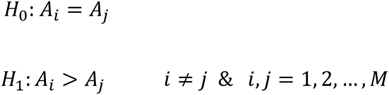

where *A_i_* and *A_j_* describe the area above the curve for immunoassays *i*, and *j*, respectively, and *j* is the total number of immunoassays tested. To test the above hypothesis, we first log-transformed immunoassay values and standardized them to a scale of 0 to 1:

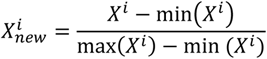

where, *X*^*i*^ is the vector of values for immunoassay *i*. The difference between the area above the curve for *i* and *j* was then calculated, denoted as *D_ij_*. We then performed a permutation test whereby we permuted for each patient sample the immunoassay identifiers and the area above the curve was recalculated for each immunoassay. This process was repeated 1,000 times (*k* = 1, 2, …, 1000). The difference between the areas above the curves for each iteration of the permutation test was defined as 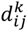. The instances that the permuted value of 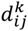 was greater than or equal to the non-permuted *D_ij_* was calculated and expressed as a fraction of the number of iterations performed, which was defined as the *P* value for testing the null hypothesis.

### Multiple linear Regression and rank-based simple linear regression

In the absence of a universal gold standard, log_10_-transformations appeared reasonable to capture immunoassay values and neutralization activity. Under our null hypothesis, a change in binding activity, log_10_(*X*), should be associated with a linear increase in neutralization, log_10_(*Y*). This relationship can be expressed as a linear regression on the log-scale:

log(*y*) = *β*_0_ + *β*_1_ log(*x*) + *∊*. A simple way of detecting departures from this model is to look for curvature in the effect of log (*x*): log(*y*) = *β*_0_ + *β*_1_ log(*x*) + *β*_2_ log(*x*)^2^ + *∊*. Any evidence that *β*_2_ is nonzero will show departure from the hypothesized relationship; for example, if higher values of binding activity produce a diminished change in neutralization efficacy, we would expect *β*_2_ to be negative. We therefore fit a multiple linear regression with the outcome variable of log-IC_50_ and each of the log-scale immunoassay variables in turn as *X*. To avoid any problematic assumptions about the distribution of the error term *∊*, the MLR was fit under a bootstrapping procedure, in which 50,000 repeated samples were taken to produce a bootstrap distribution of the parameter estimates. This was used to compute non-parametric 95% confidence intervals for the *β*_2_ quadratic effects.

In addition to this MLR approach using log-transformed assay values, we conducted a series of rank-based analyses. Rather than focusing on the ratio-relationship directly, we hypothesized that high neutralization values (relative to the sampling distribution) should correspond to high binding values (relative to the sampling distribution), in such a way that the rank-ratios, 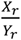, should follow a distribution with mean not depending on the binding rank, *X_r_*. This was investigated via a bootstrapped simple linear regression with the rank-ratio of binding to neutralization as the outcome, and the binding rank as the single explanatory variable. Under the null hypothesis, the slope parameter for the binding rank, *β*_1_, should be equal to zero. We again performed 50,000 repeated samples to produce bootstrap distributions and corresponding non-parametric confidence intervals for *β*_1_.

## ACKNOWLEDGEMENTS

We thank all blood donors that contributed samples to this study and Julie Kurt from the Department of Pathology and Drs. Mary Rysavy and Kim Kenne from the Department of Gynecology and Obstetrics at the University of Iowa for assistance in coordinating these studies. We also thank Michelle Sexton of the Iowa State Hygienic Laboratory and Dr. Louis Katz of the Mississippi Valley Regional Blood Center for assistance in conducting the immunoassays. This work was supported by the Department of Pathology at the University of Iowa. DWB was supported by NIH T32 AI007511. KR and HVE were supported by NIH T32GM007337. HVE and NR were supported by NIH R01AI134733 and R21 AI144215 to WJM. The funders had no role in study design, data collection and interpretation, or the decision to submit the work for publication. All corresponding authors had full access to all the data in the study and had final responsibility for the decision to submit the manuscript for publication.

## SUPPLEMENTAL FILES

**Supplemental Figure S1.**
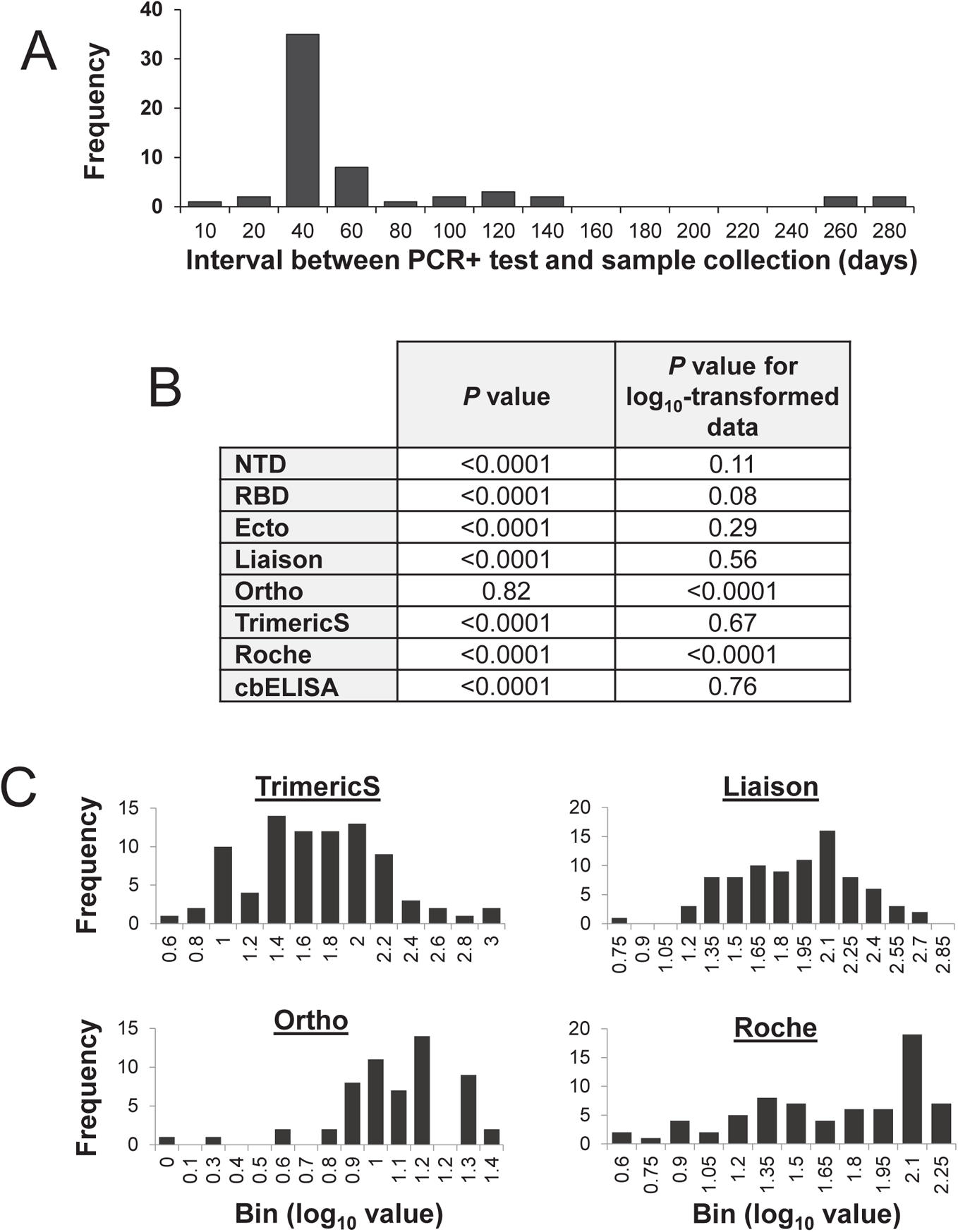
Measurements of spike-targeting antibodies in 85 convalescent samples using immunoassays that apply different components of SARS-CoV-2 as capture antigens. **(A)** Distribution of the time interval between the PCR test indicating infection by SARS-CoV-2 and collection time of the samples used for this study. **(B)** Normality tests of immunoassay values. The Shapiro-Wilk test was performed for the immunoassay values and for the log_10_-transformed immunoassay values. The null hypothesis for this test is that the data are normally distributed. *P* values lower than 0.05 indicate that the null hypothesis is rejected. **(C)** Distribution of immunoassay values measured for 85 convalescent using commercial tests.

**Supplemental Figure S2.**
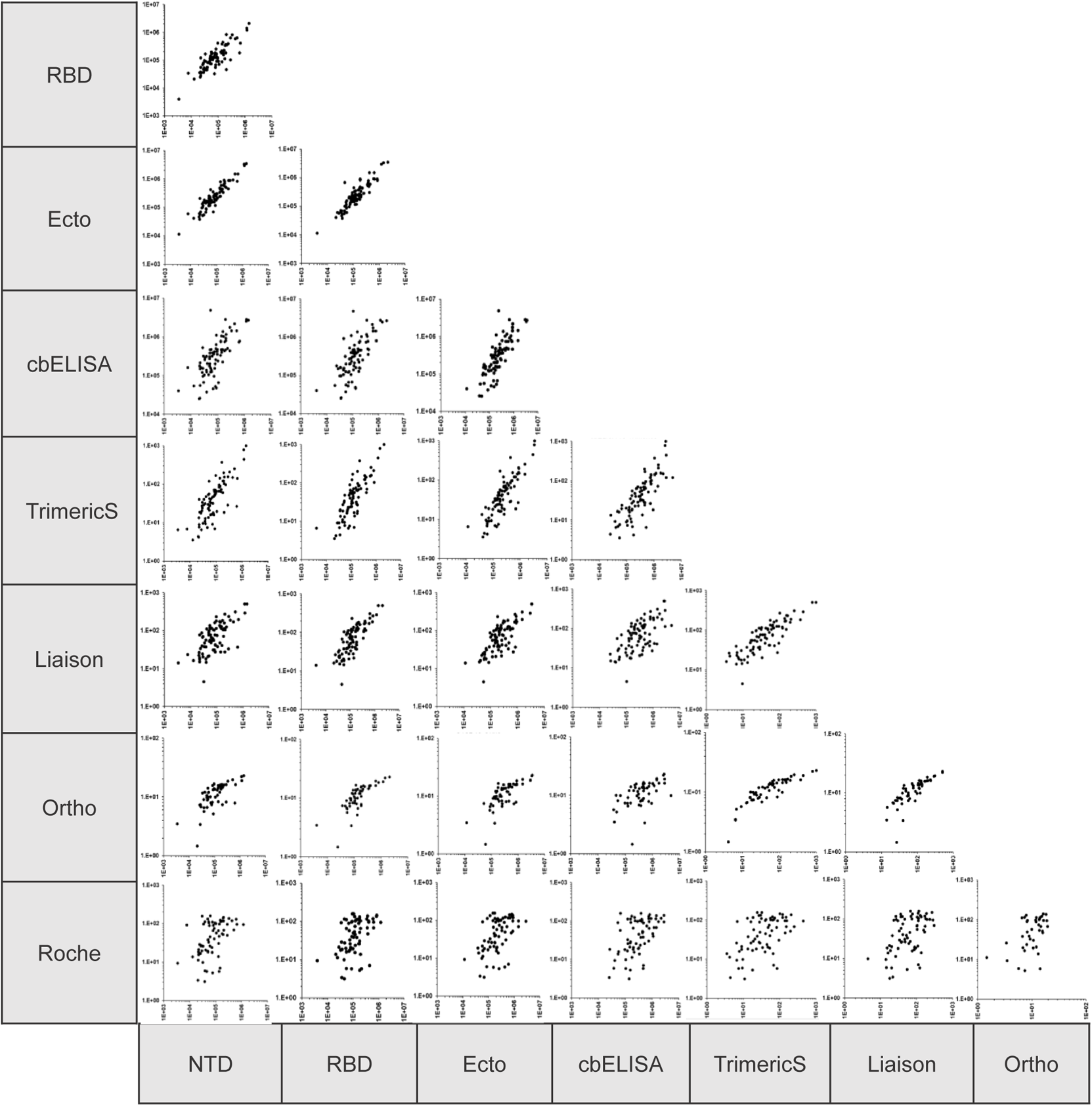
Comparison of the log_10_-transformed values measured in eight immunoassays for 85 convalescent serum and plasma samples. Values in the commercial assays are expressed in arbitrary units (AUs) whereas values in ELISAs are expressed in relative light units (RLUs). Values in the x and y axes are shown in log_10_ scale. Correlation coefficients are shown in **Fig. 1C**.

**Supplemental Figure S3.**
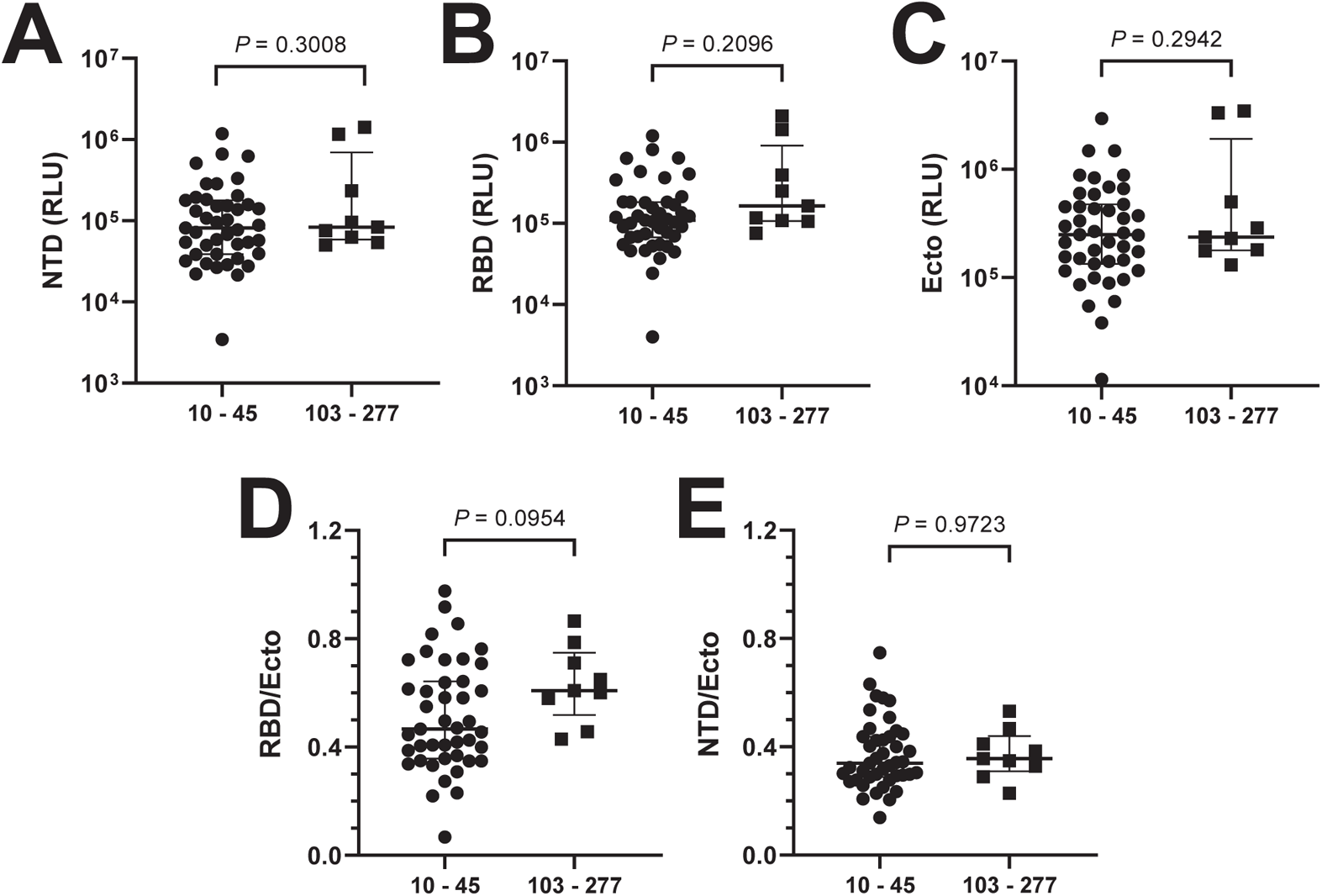
Relationship between timing of sample collection and target specificity of the antibody response. The interval (in days) between the PCR-positive test indicating SARS-CoV-2 infection and the time of plasma or serum collection for these studies was determined. **(A-C)** Immunoassay values are compared between samples with an interval of 10-45 days (n=43), and samples with an interval of 103-277 days (n=9). **(D,E)** Comparison of the RBD/Ecto or NTD/Ecto ratios for samples collected after the indicated intervals from detection of infection by PCR. *P* Value, two-tailed test.

**Supplemental Figure S4.**
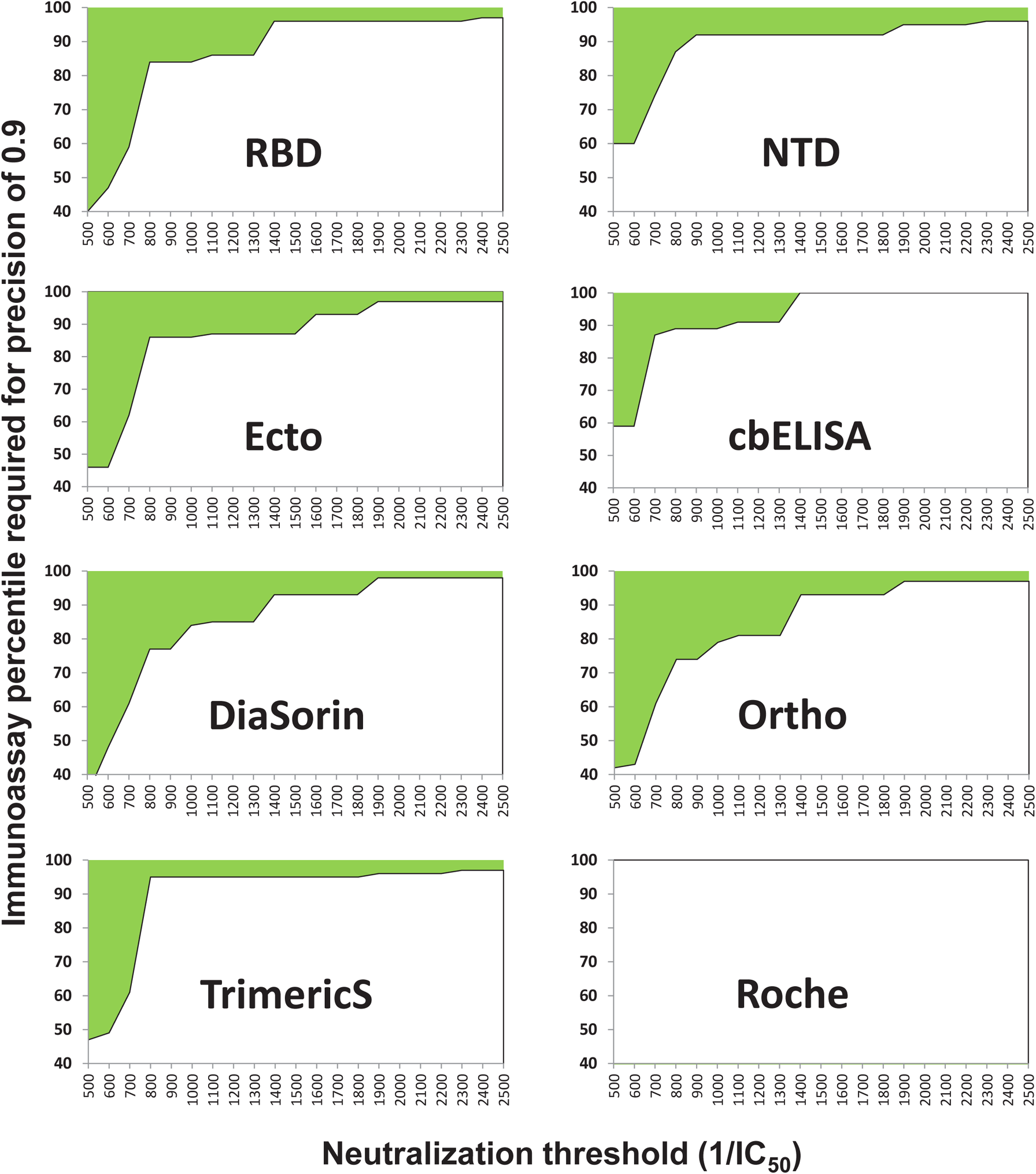
Immunoassay percentiles required to predict neutralization at the indicated thresholds with a precision of 0.9. The shaded area describes the combination between neutralization thresholds and sample immunoassay percentiles that allow prediction with a precision of 0.9 or higher. The Roche test did not achieve a precision of 0.9, and thus a value could not be computed for this assay.

**Supplemental Figure S5.**
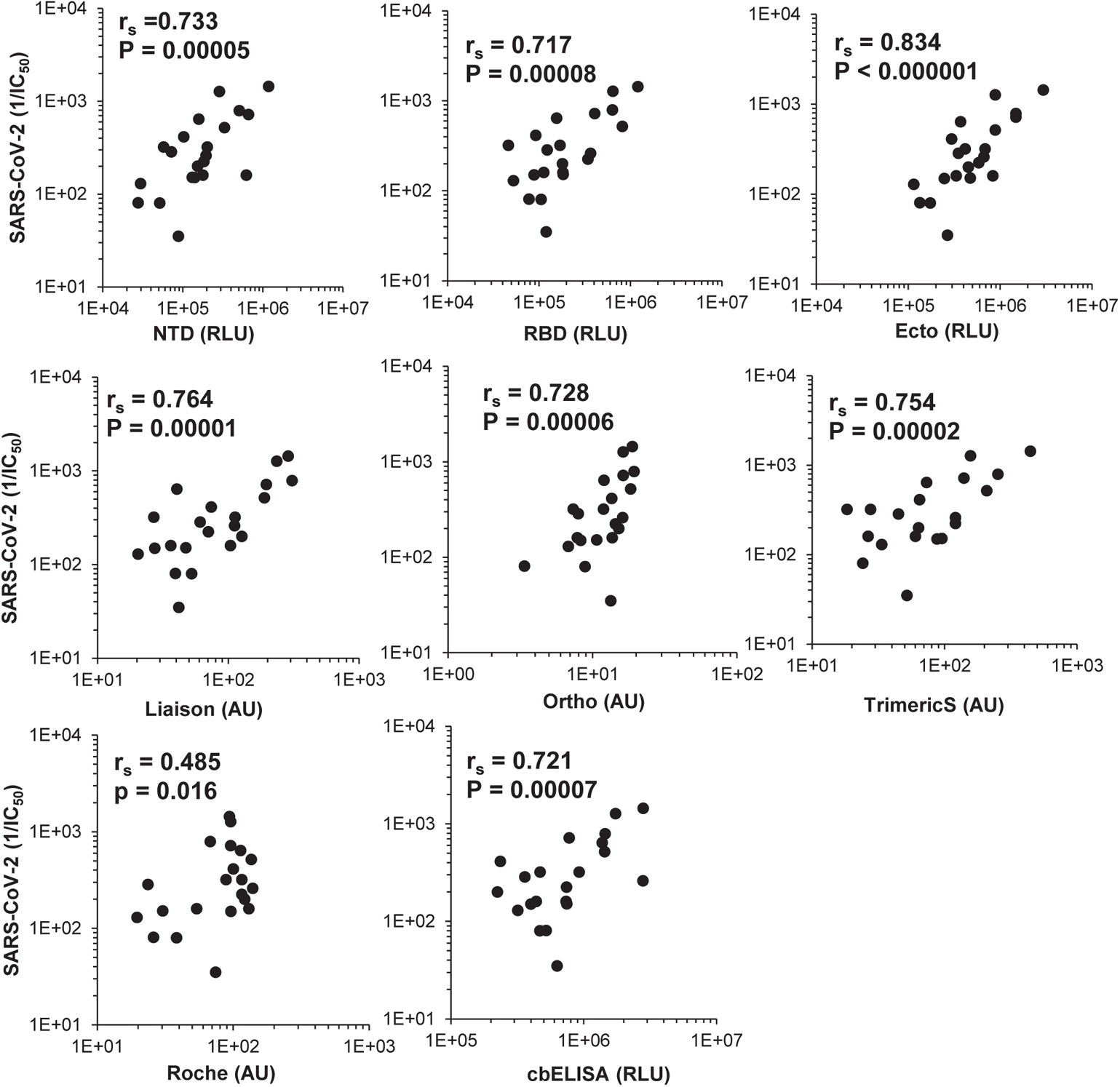
Relationships between immunoassay values of COVID-19 convalescent samples and their neutralization of replicative SARS-CoV-2. r_S_, Spearman correlation coefficient. *P* value, two-tailed test.

**Supplemental Figure S6.**
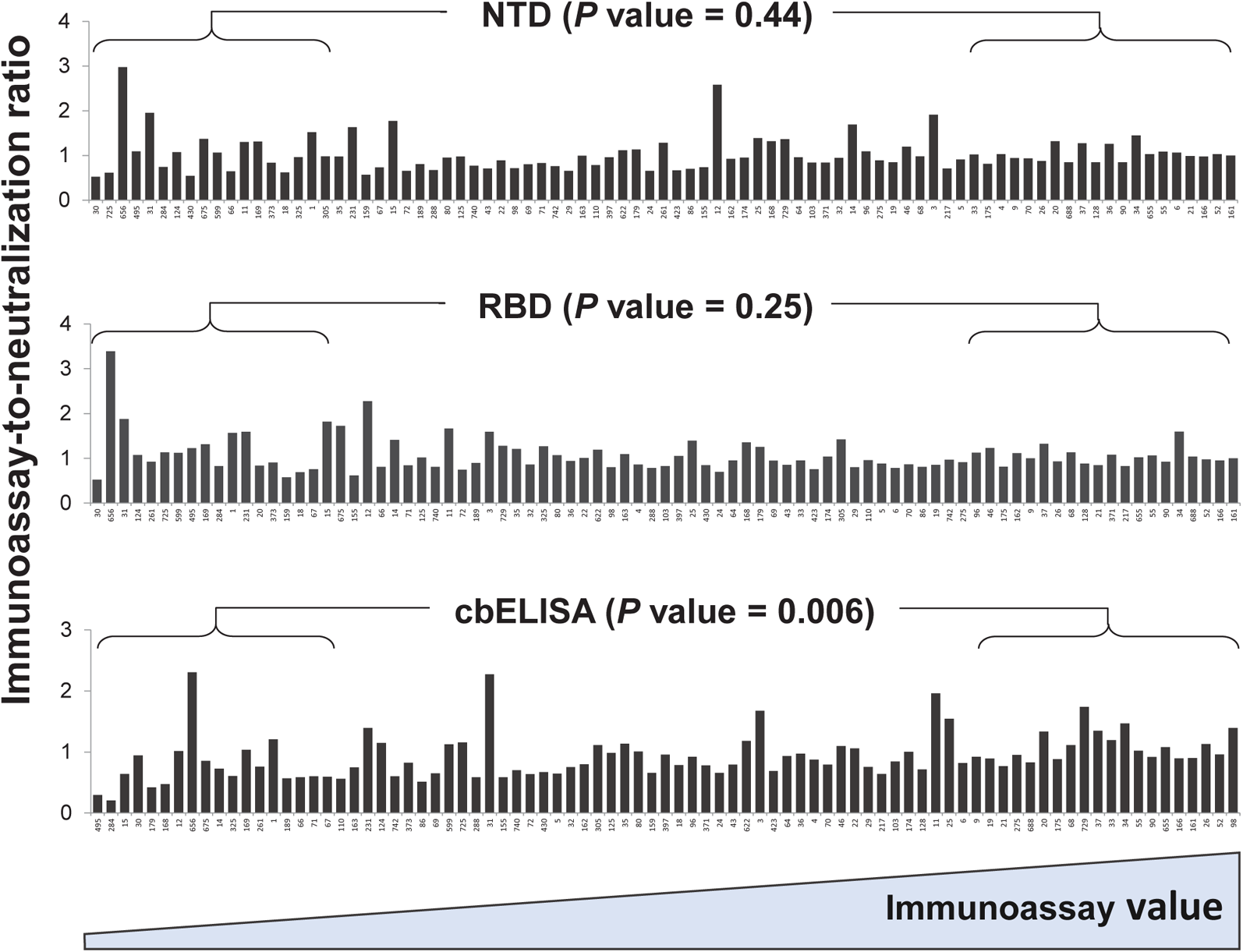
Relationship between the level of spike-specific antibodies in convalescent samples and their relative neutralization potency. IC_50_ values of convalescent samples and their immunoassay values were log-transformed and adjusted to a scale of 0.1 to 1. For each sample, the ratio between the immunoassay value and the IC_50_ value was calculated and shown. Samples are ordered by increasing immunoassay values from left to right. The ratios calculated for the 20 samples with the lowest immunoassay values and the 20 samples with the highest immunoassay values were compared using an unpaired T test; the *P* values for a two-tailed test are indicated.

**Supplemental Table S1.** Sample donor information. n.a., indicated data not available.

